# The gut mycobiome of healthy mice is shaped by the environment and shapes metabolic outcomes in response to diet

**DOI:** 10.1101/2020.06.25.158287

**Authors:** Tahliyah S. Mims, Qusai Al Abdullah, Justin D. Stewart, Sydney P. Watts, Catrina T. White, Thomas V. Rousselle, Amandeep Bajwa, Joan C. Han, Kent A. Willis, Joseph F. Pierre

## Abstract

**Objective:** As an active interface between the host and their diet, the gut bacteriome influences host metabolic adaptation. However, the contribution of gut fungi to host metabolic outcomes is not yet understood. Therefore, we aimed to determine if host metabolic response to an ultra-processed diet reflects gut fungal community composition.

**Design:** We compared jejunal fungi and bacteria from 72 healthy mice with the same genetic background but different starting mycobiomes before and after 8 weeks on an ultra-processed or standardized diet using 16S and internal transcribed spacer region 2 ribosomal RNA sequencing. We measured host body composition using magnetic resonance imaging, examined changes in metabolically active host tissues and quantified serum metabolic biomarkers.

**Results:** Gut fungal communities are highly variable between mice, differing by vendor, age and sex. After exposure to an ultra-processed diet for 8 weeks, persistent differences in fungal community composition strongly associate with differential deposition of body mass in male mice compared to mice on standardized diet. Fat deposition in the liver, genomic adaptation of metabolically active tissues and serum metabolic biomarkers are correlated with alterations in fungal diversity and community composition. Variation in fungi from the genera *Thermomyces* and *Saccharomyces* most strongly associate with increased weight gain.

**Conclusions:** In the gut of healthy mice, host-microbe metabolic interactions strongly reflect variability in fungal communities. Our results confirm the importance of luminal fungal communities to host metabolic adaptation to dietary exposure. Gut fungal communities may represent a therapeutic target for the prevention and treatment of metabolic disease.

**Graphical Abstract:** 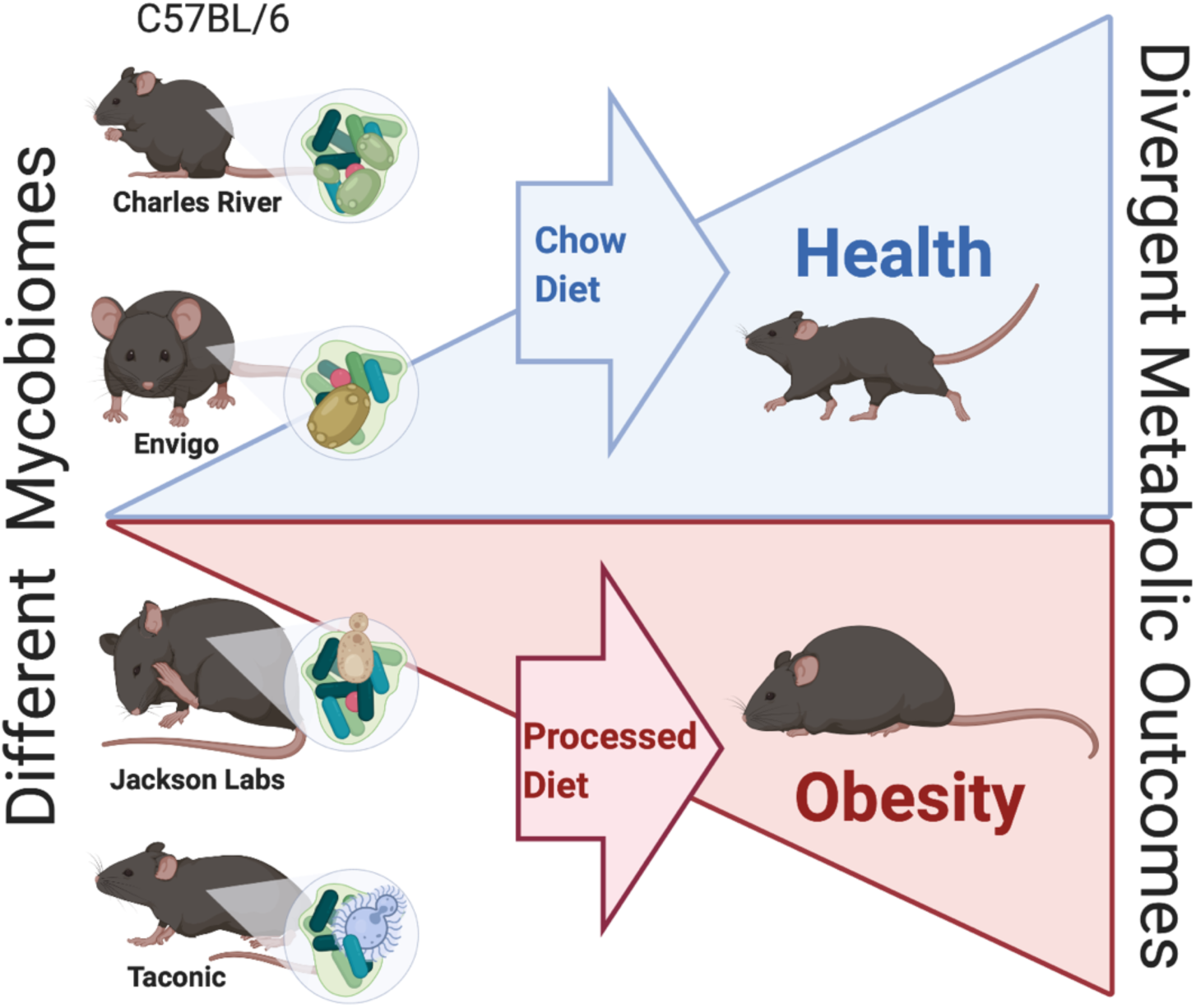

**In Brief:**

- What is already known about this subject? Gut bacterial communities have evolved to influence the metabolic outcomes of the host in mammals. Evidence from across the lifespan suggests that differences in composition of these communities is associated with energy consumption. However, gut microbial communities, while often equated to bacteria, are diverse, multi-kingdom ecologies and limited information is available for the role of other kingdoms of life, such as fungi.
- What are the new findings? Gut fungal communities, collectively termed the mycobiome, are less diverse and abundant than bacterial communities in the gastrointestinal tract. This study identifies the considerable influence of the environment and dietary exposure on the composition of jejunal fungal communities in healthy mice with the same genetic background. After exposure to processed diet, differences in fungal community composition in male mice were strongly correlated with persistent differences body composition and markers of metabolic tone.
- How might it impact on clinical practice in the foreseeable future? These results verify that the baseline metabolic tone of health mice strongly reflects the ecological complexity of the gastrointestinal mycobiome. Variation in the composition of gut fungal communities is likely an underappreciated source of experimental and clinical variability in metabolic studies. Gastrointestinal fungi are likely a target for prevention and treatment of metabolic disease.

## INTRODUCTION

The modern diet is dominated by ultra-processed sugar and carbohydrates that are linked to metabolic and immune-mediated diseases.^1^ Disruption of the gut microbiome also influences the development of metabolic disease,^2,3^ and dietary composition is a key driver of gut microbial community structure and function.^1^ Gut microbes form an interface between the diet and the host, participate in digestion and energy extraction from otherwise indigestible fiber and oligosaccharides,^4^ produce short-chain fatty acids and novel metabolites, and ultimately shape host endocrine and immune signaling. Given these findings, robust microbial communities appear to be a crucial component of metabolic homeostasis as supported by data across diverse species, life stages, and disease states.^2,5^

While the gut microbiome is often equated to bacteria, microbial communities contain diverse populations of archaea, viruses, protists and fungi.^6^ Collectively termed the mycobiome, gut fungal communities of molds and yeasts are crucial to maintaining gut homeostasis and systemic immunity.^6^ Mounting evidence suggests other kingdoms of life, such as viruses, can also influence host metabolic tone.^7^ However, data describing the role of the mycobiome in host metabolism are scarce.^8,9^ Recent studies in humans and mice indicate commensal fungi have the potential to influence host metabolism directly^6,10-13^ and via alterations to bacterial community composition.^14,15^

Therefore, whether the gut mycobiome influences host metabolism remains unclear. This question is further obscured by the difficulty of sorting environmental and transiently ingested fungi from true colonizers, especially in mobile human populations. Thus, we designed a study using specific pathogen-free mice with the same genetic background so that we could control for age, sex and previous founder exposures. We also sourced from four different vendors to vary the initial mycobiomes. Utilizing this design, we tested our hypothesis that the gut mycobiome shaped the host metabolic response to an ultra-processed diet.

We found the gut mycobiome of laboratory mice differs dramatically between animal vendors. Using a combinatory approach, we identified fungal populations that shifted in response to processed diet and identified key fungal species that may be linked to host metabolic alterations. We show baseline correlations in the gut mycobiome strongly associate with changes in host adiposity and serum metabolic biomarkers in response to highly processed low-fat diet. Our results support the role of the gut mycobiome in host metabolic adaptation and have important implications regarding the interpretation of microbiome studies and the reproducibility of experimental studies of host metabolism.

## METHODS

Detailed methods are included in the online supplementary materials.

### Ethics Statement

All animal studies were approved by the Institutional Animal Care and Use Committee at the University of Tennessee Health Science Center. Laboratory animal care policies follow the Public Health Service Policy on Humane Care and Use of Laboratory Animals. All results are reported in a manner consistent with ARRIVE guidelines.^16^

### Mice

Eight-week-old specific pathogen-free C57Bl/6 mice of both sexes were purchased from Charles River Laboratories (CR, Wilmington, MA), Envigo (ENV, Indianapolis, IN), Taconic Biosciences (TAC, Rensselaer, NY) and Jackson Laboratories (JAX, Bar Harbor, ME). Mice were maintained in microisolators to prevent contamination. On arrival, 6 mice from each vendor were humanely euthanized, and jejunal and fecal contents were collected to quantify the baseline mycobiome. The remaining 12 mice/group were randomized to chow (Envigo 7912) or purified processed (Research Diets Inc. D12450B, New Brunswick, NJ) diet and monitored biweekly for 8 weeks to quantify food intake and body composition. Fecal and jejunal contents were collected to quantify the microbial communities at the end of the experiment. Details regarding exposure, housing, tissue collection and sample processing are detailed in the online supplement.

### Fungal and Bacterial Ribosomal RNA Extraction

Microbial DNA was extracted using lyticase and proteinase K, amplified and sequenced according to our previously published protocols.^17^ Sequencing was performed using the MiSeq platform (Illumina, San Diego, CA) for both the 16S and internal transcribed spacer region 2 (ITS2) ribosomal (r)DNA genes at the Argonne National Laboratory (Lemont, IL).

### Body Composition

Body composition (fat and lean mass) were measured biweekly using an EchoMRI 1100 system (EchoMRI, Houston, TX). These differences were confirmed by tissue collection after the experiment ended.

### Serum Metabolic Biomarkers

Serum triglycerides were quantified using a Triglyceride Colorimetric Assay Kit (Cayman Chemicals, Ann Arbor, MI). Metabolically active hormones were quantified using the Bio-Plex Pro Mouse Diabetes Panel (Bio-Rad Laboratories, Hercules, CA). Both were prepared according to the manufacturer’s instructions. Details are presented in the online supplement.

### Histochemistry

Liver histology using hematoxylin and eosin was performed on paraldehyde-preserved tissue (Vector Laboratories, Burlington, Canada). Details are presented in the online supplement.

### Gene Expression

Gene expression in liver and epididymal white adipose tissue was quantified using SYBR Green qRT-PCR (Applied Biosystems, Inc., Foster City, CA) on the QuantStudio 6 Flex Real-Time PCR System (Applied Biosystems). Procedural details and gene targets are described in the online supplement.

### Statistical Analysis

Sequence data were processed in QIIME 1.9 and analyzed in Calypso 8.84^18^ (Hellinger transformed) and R 3.6.0 primarily using the *vegan* and *mvabund* (model-based analysis of multivariate abundance data) packages.^19^ Biochemical data were analyzed using R. Machine learning was performed in R and presented in accordance with transparent reporting of a multivariable prediction model for individual prognosis or diagnosis (TRIPOD) guidelines.^20^ Details of the bioinformatics and statistical analyses are presented in the online supplement.

## RESULTS

To determine if the gut mycobiome altered host metabolism, we studied 72 mice obtained in one shipment each from four vendors (CR, ENV, JAX and TAC). We characterized the fungal and bacterial communities of the jejunum, the most diverse fungal populations in the mouse gut.^9^ We quantified fungal communities using ITS2 rDNA gene sequencing, the most reliable culture-independent technique for detecting mammalian-associated fungi.^21^ We obtained 960,043 total ITS2 reads, with an average of 14,329 reads per sample. We detected no significant environmental or procedural contamination in jejunal samples. *Fusarium* was the most frequent potential contaminate detected (**Supplemental Figure 1**).

### Interkingdom Composition of Murine Diets

A fundamental question of gut mycobiome research is the extent to which fungal organisms detected by next generation sequencing represent organisms present in the diet or true commensal colonization by organisms replicating within the gut lumen. Therefore, we sequenced the food pellets that arrived with the mice from their respective vendors as well as the standardized chow and processed diets that the mice were fed in our experimental isolators. On arrival, diets from the vendors showed similar multi-kingdom microbial composition (**Supplemental Figure 2**), distinct from the experimental diets and different from microbes in the guts of the mice themselves.

### Gut Fungal Communities Cluster by Vendor

The gut bacterial communities of experimental mice differ markedly by vendor.^22^ This variability has been utilized to understand the interrelatedness of gut and lung microbiota^23^ and standardize microbiota for genetic studies.^24^ We sought to test to what extent the vendor-induced variability in gut bacterial communities held true for gut fungal communities. Upon arrival from the vendor, the gut fungal communities of healthy laboratory mice from each vendor were distinct (p =0.001, R^2^ =0.211, permutational multivariate analysis of variance, PERMANOVA; p =0.575, permutational multivariate analysis of dispersion, PERMDISP2), with unique composition (p =0.001, *f* =1.1; canonical correspondence analysis, CCA), despite similar diversity (p =0.53, f =0.77; ANOVA Chao1 Index, **Figure 1**). Across the four vendors, a core mycobiome of 18 genera was identified. An additional 14 genera were identified in animals from multiple venders. Mice from JAX hosted the most unique genera (5: *Rasamsonia, Mycosphaerella, Millerozyma, Geosmithia* and *Byssochlamys*) followed closely by TAC (*Bipolaris, Hanseniaspora* and *Tritirachium*), then CR Laboratories (*Lichtheimia* and *Dioszegia*), while only one genus (*Plectosphaerella*) was unique to animals from ENV (**Supplemental Figure 3**).

**Figure 1.**
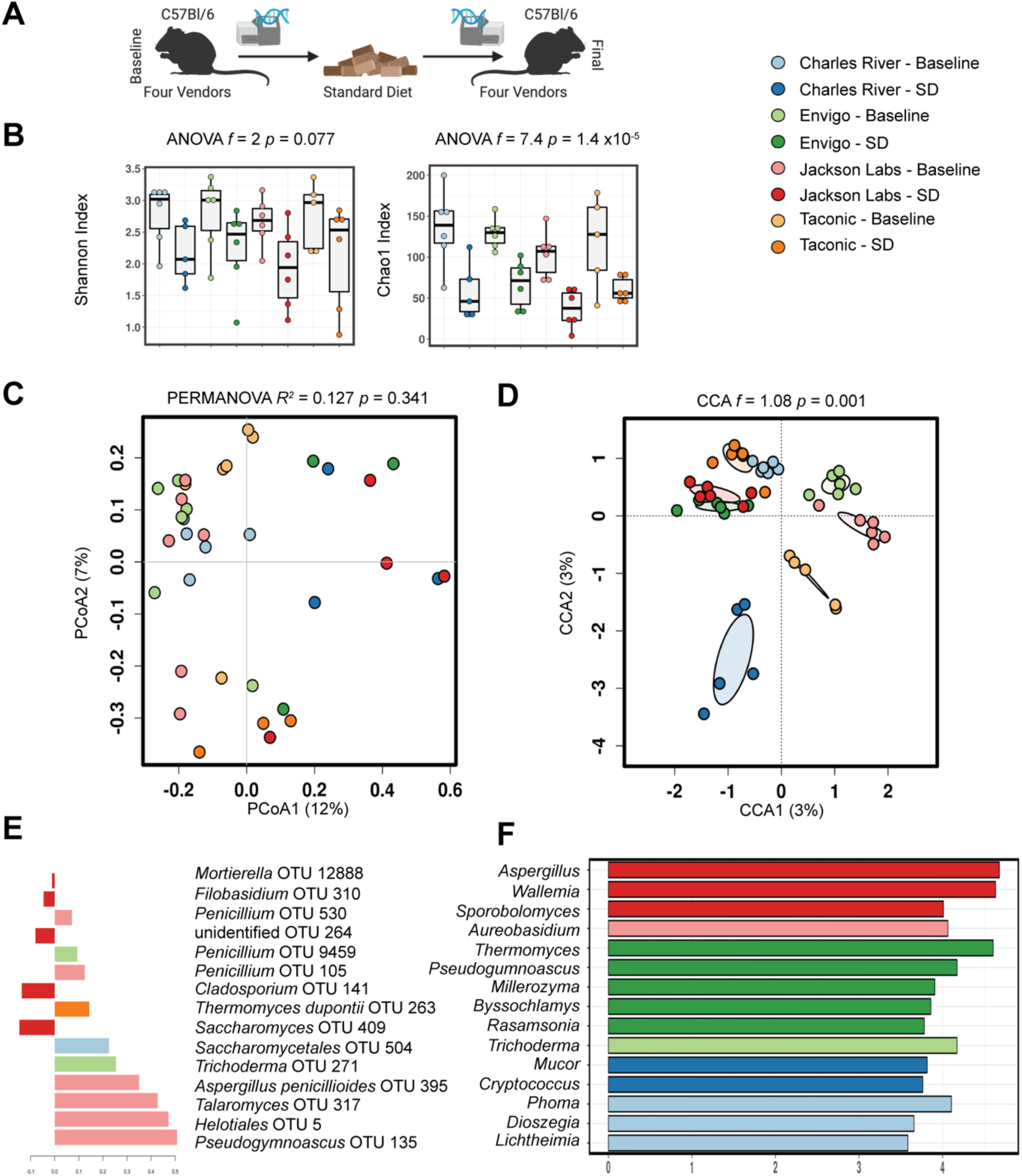
Gut fungal communities cluster by vendor, age and dietary exposure. (*A*) Experimental schematic. (B) Compared to healthy mice exposed to standardized chow diet for 8 weeks, baseline community diversity is higher in mice upon delivery from vendors. While gut fungal communities remained distinctly clustered by vendor, exposure to standardized chow diet for 8 weeks exerted a convergent effect on community composition (*C* and *D*, Bray-Curtis dissimilarity distance). Supervised partial least squares discriminant analysis (*E*) and linear discriminant analysis of effect size (*F*) confirm key operational taxonomic units and genera driving differences in community composition. Hypothesis testing was performed using ANOVA (*B*), PERMANOVA (*C*) and CCA (*D*). CCA, canonical correspondence analysis; OTUs, operational taxonomic units; SD, standard diet; PERMANOVA, permutational multivariate ANOVA; PCoA, Principal coordinates analysis. Schematic illustrated using BioRender.

### Gut Fungal Community Diversity Declines with Age and Exposure to Standard Diet

While highly variable, complexity of the gut bacteriome declines with maturity^25-27^, a finding we confirmed (**Supplemental Figure 7**). Data on longitudinal trends in gut mycobiome development are lacking. After we determined gut fungal communities in mice differed by their initial environment, we asked how these communities evolved in a different environment. To determine the temporal dynamics of the gut mycobiome, we exposed mice with the same genetic background from different vendors to the same standardized diet and microenvironment for 8 weeks (**Figure 1A**). Community diversity declined in all animals exposed to standardized diet (p =1.4 × 10^−5^, f =7.4; Chao1 Index, **Figure 1B**). Community composition similarly converged with time, resulting in more similar, less complex, but still distinct fungal communities (p =0.341, R^2^ =0.127; PERMANOVA. p =0.215; PERMDISP2, **Figure 1C**. p =0.001, f =1.08; CCA, **Figure 1D**). Supervised partial least squares discriminant analysis confirmed key operational taxonomic units driving differences in community composition primarily result from loss of unique taxa (**Figure 1E**).

In general, fungal community differences between the vendors converged, resulting in communities with overall similar diversity and more similar community composition. A reduced core of 12 shared genera remained, with animals from CR and TAC being the most divergent (**Supplemental Figure 5**). To further understand the temporal dynamics of fungal communities, we built a series of generalized linear models using *mvabund* that revealed fungal and bacteria communities differ by diet and timepoint (**Table 1, Supplemental Figure 6**). Community diversity decreased with time, leading to 5 taxa being selected against and only one taxon, *Aspergillus*, persisting. In contrast, bacterial communities differed by vendor and not sex. From these analyses, we conclude jejunal fungal communities are more dynamic and susceptible to environmental influences than bacterial communities.

**Table 1.** Determinants of microbial community composition in healthy mice.

| Variable | Fungal |  | Bacterial |  |
| --- | --- | --- | --- | --- |
|  | Likelihood Ratio Test<sup>†</sup> | P Value<sup>‡</sup> | Likelihood Ratio Test<sup>†</sup> | P Value<sup>‡</sup> |
| Vendor | 237.9 | 0.153 | 874.5 | 0.032 |
| Diet | 132.5 | 0.004 | 843.5 | 0.001 |
| Sex | 159.5 | 0.036 | 288.8 | 0.756 |
| Timepoint | 252.3 | 0.001 | 333.3 | 0.007 |

<sup>†</sup>The likelihood ratio test quantifies the goodness-of-fit between generalized linear models (*mvabund*).
<sup>‡</sup>P-values were estimated using parametric bootstrapping with 999 permutations.

### Gut Fungal Community Composition Evolves on Exposure to Processed Diet

We demonstrated that environmental differences between gut fungal communities converged after exposure to a standardized diet and environment (**Figure 1**). Given that gut bacterial communities are influenced by diet,^28^ we sought to understand how the mycobiome was shaped by exposure to an ultra-processed diet. To address this question, we exposed a second group of mice from the same vendors to a processed diet for 8 weeks. Compared to baseline, the community diversity of fungal communities declined after exposure to processed diet (p =0.0038, f =4; ANOVA, Chao1 Index, **Figure 2A**). The community composition was also markedly different (p =0.00033, R^2^ =0.278; PERMANOVA, **Figure 2B**. p =0.001, R =0.347; analysis of similarities (ANOSIM), **Figure 2C**, p =0.004, *mvabund*, **Table 1**). This change resulted in a convergence of microbial communities that were no longer distinct by PERMANOVA and had similar overall diversity (**Supplemental Figure 8, Table 1**). Nonetheless, more sensitive generalized linear modeling identified significant differences before and after exposure to a processed diet (p =0.001, *mvabund*, **Table 1**). Interkingdom community structure showed a similar pattern (**Supplemental Figure 9**).

**Figure 2.**
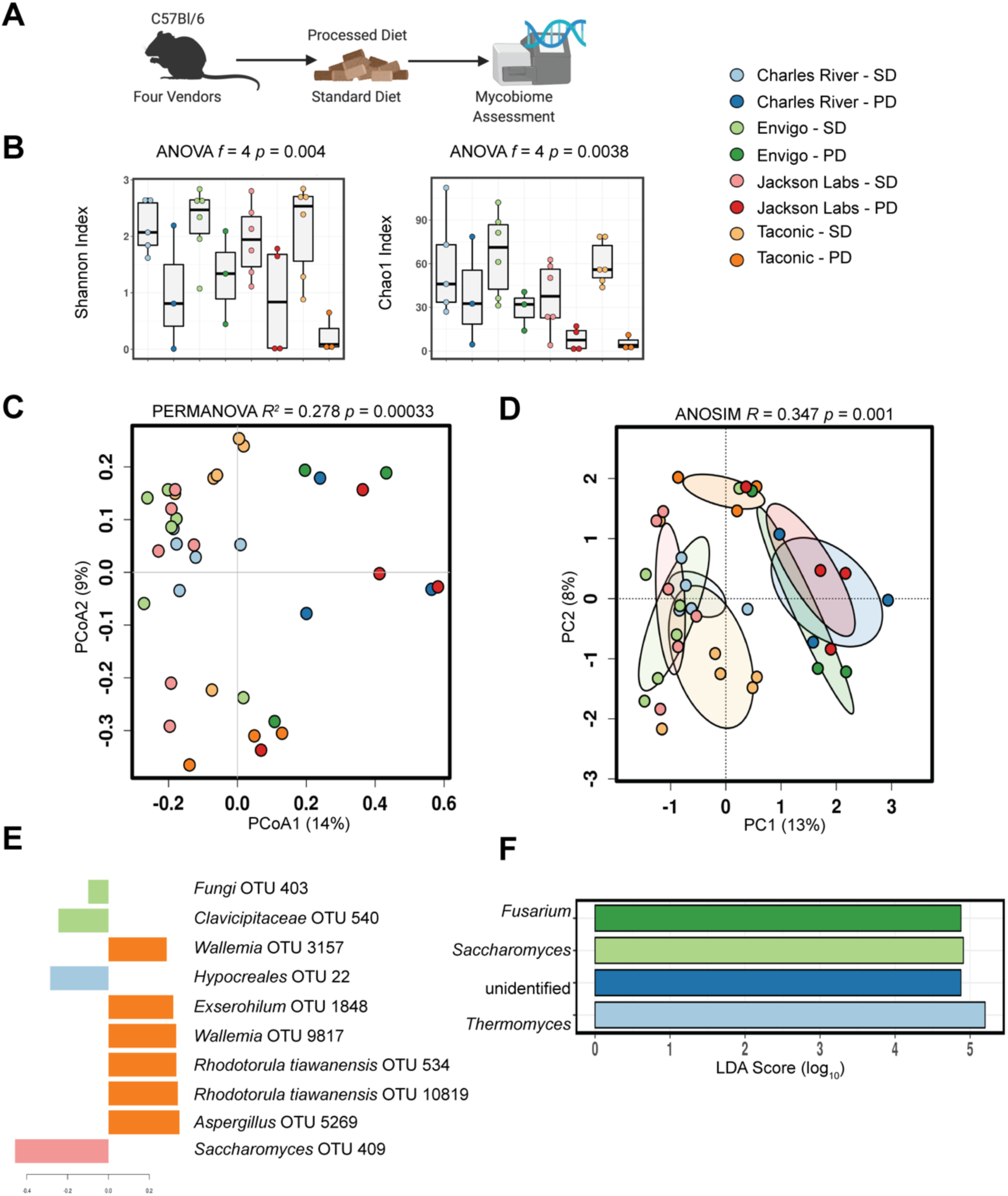
Exposure to processed diet produces a more pronounced alteration of gut fungal communities than does exposure to standardized chow. (*A*) Experimental Schematic. (*B*) Compared to healthy mice exposed to standardized chow diet for 8 weeks, mice exposed to processed diet show reduced community diversity. While gut fungal communities remained distinctly clustered by vendor, exposure to processed diet for 8 weeks exerted a convergent effect on community composition that exceeded the similar effect of standardized chow (*C* and *D*, Bray-Curtis dissimilarity distance). Supervised partial least squares discriminant analysis (*E*) and linear discriminant analysis of effect size (*F*) confirm key operational taxonomic units and genera driving differences in community composition. Hypothesis testing was performed using ANOVA (*B*), PERMANOVA (*C*) and ANOSIM (*D*). ANOSIM, analysis of similarities; CCA, canonical correspondence analysis; OTUs, operational taxonomic units; LDA, linear discriminant analysis; PERMANOVA, permutational multivariate ANOVA; PD, processed diet; PCA, principal components analysis; PCoA, principal coordinates analysis; SD, standard diet. Schematic illustrated using BioRender.

### Gut Fungal Community Composition Differs Between Standard and Processed Diet

We next asked how the fungal communities in mice exposed to processed diet differed from mice exposed to standardized chow diet. While a similar overall trend in reduction in diversity and separation of multivariate clustering can be seen in all animals, differences between the community composition of each vendor remained (**Figure 3**). The diversity of animals from TAC and JAX was the most reduced, with CR having the most retained diversity. The clear divergence of processed diet exposed animals is also apparent on multivariate analyses (p =0.01, f =1.08; CCA, **Figure 3**). The shared core microbiome previously observed at baseline or after exposure to standardized chow collapsed with only one genus, *Aspergillus*, shared between all groups. Animals from CR had the most retained diversity, with the noteworthy unique prevalence of the genera *Candida* and *Aureobasidium* (**Supplemental Figure 8**).

**Figure 3.**
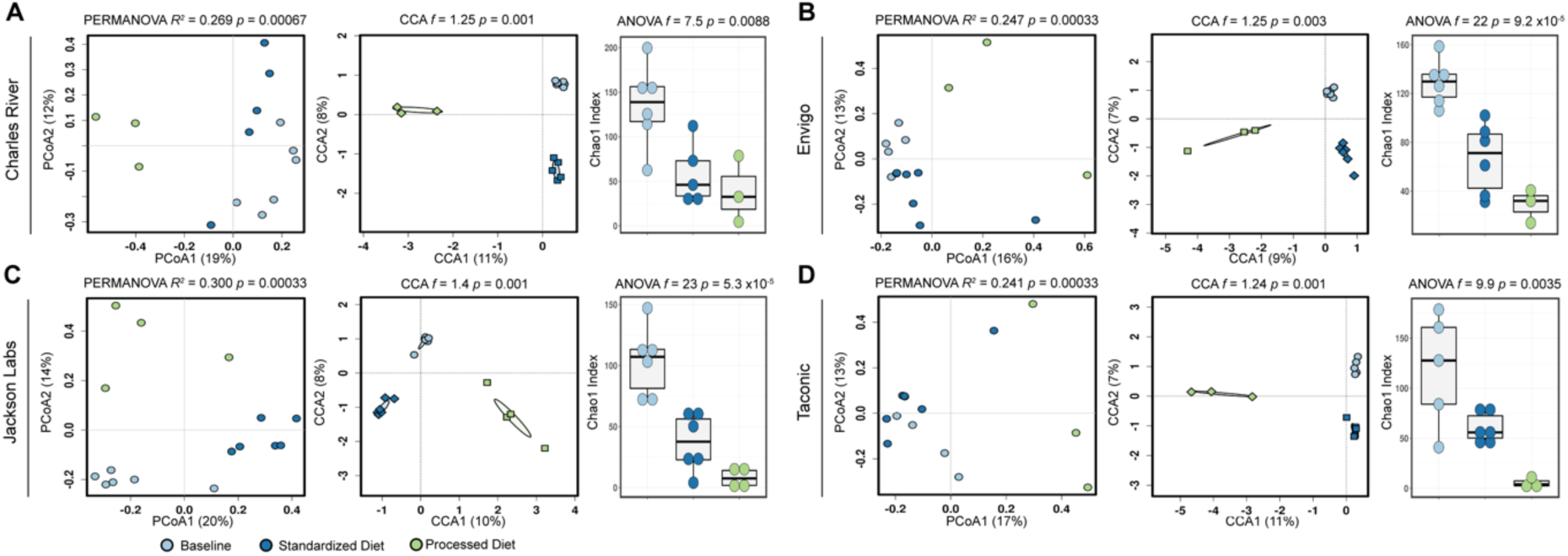
Baseline fungal composition influences final community composition. Differences in fungal community composition and diversity present on arrival from the four laboratory mouse vendors lead to persistent differences in the composition and diversity of fungal communities (*A*-*D*). Hypothesis testing for community composition was performed using permutational multivariate ANOVA and canonical correspondence analysis of Hellinger transformed Bray-Curtis dissimilarity distances and community diversity by ANOVA of the Chao1 index. CCA, canonical correspondence analysis; OTUs, operational taxonomic units; PERMANOVA, permutational multivariate ANOVA; PCoA, Principal coordinates analysis.

### Differences in Fungal Community Composition Persist Despite Exposure to Processed Diet

To identify differences between vendors after exposure to processed diet versus standardized chow, we utilized a series of complementary approaches, including analysis of composition of microbiomes (ANCOM), mixed effect regression and discriminant analysis of principal components. In animals from JAX, the most robust changes were a reduction in the genus *Saccharomyces* and corresponding bloom in *Thermomyces*, while animals from ENV demonstrated only the reduction in *Saccharomyces* and animals from CR the bloom in *Thermomyces*. Animals from TAC showed the opposite effect: *Thermomyces*, which was already at low levels in animals exposed to standard diet, was eliminated after exposure to processed diet, as was the genus *Penicillium* (**Figure 4**). Furthermore, generalized linear modeling identified three specific taxa across taxonomic resolution that were associated with exposure to the processed diet: the orders Helotiales (p <0.001) and Saccharomycetales (p =0.049) and the genus *Wallemia* (p =0.014, **Supplemental Figure 6**).

**Figure 4.**
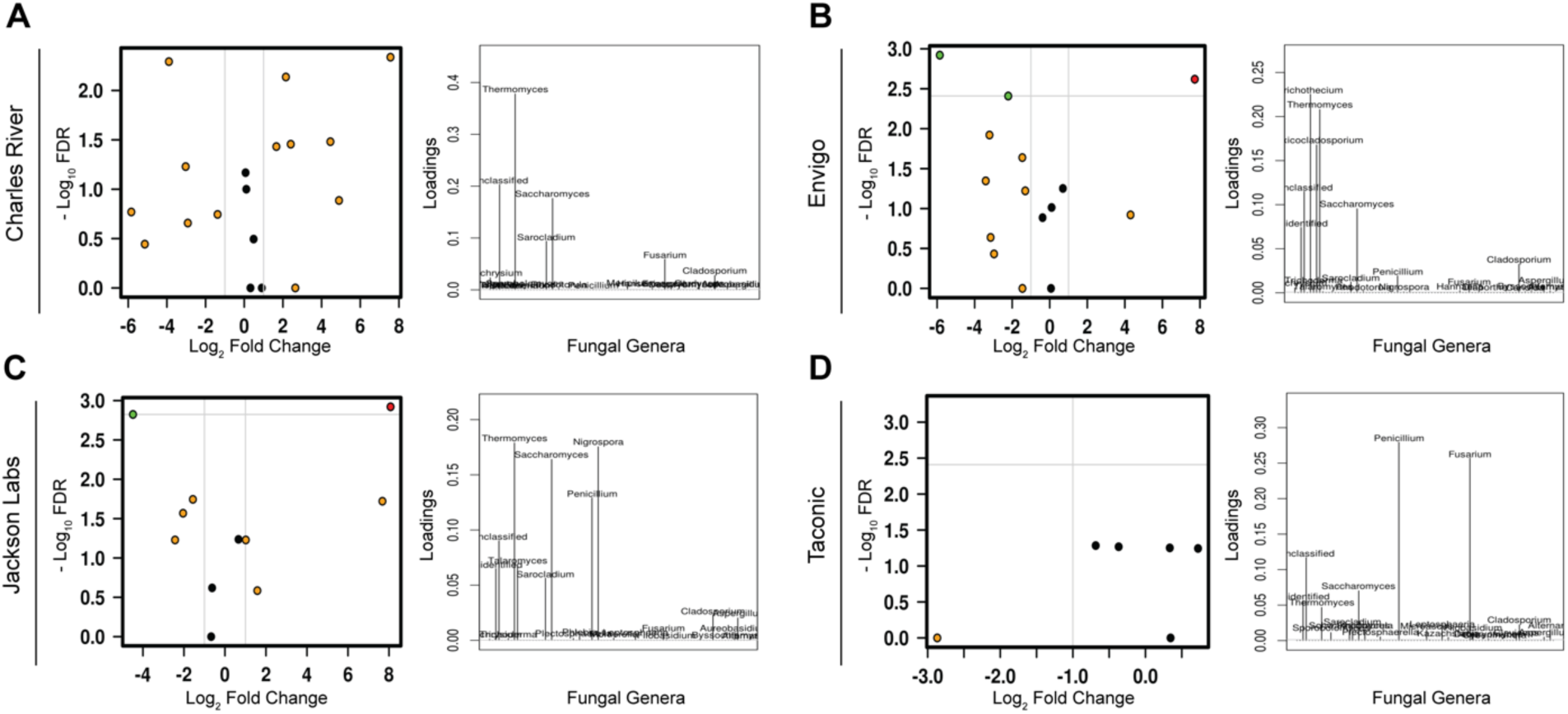
Differences among animals sourced from different vendors persist after exposure to processed diet. Volcano plots of mixed-effect regression identify differences between animals exposed to processed diet that are highlighted by loading plots of discriminant analysis of principal components (*A-D*). FDR, false discovery rate.

### Exposure to Processed Diet Reduces Fungi-Associated Co-Occurrence Networks

While overall interkingdom co-occurrence patterns among taxa did not significantly differ after exposure to the processed diet (p =0.3438, t-test; **Figure 5**), there was a noticeable reduction in the number and magnitude of co-occurrent relationships. This reduction was largely attributed to changes in 4 fungal taxa (Helotiales, *Trichothecium, Toxicocladosporium* and *Filobasidium*).

**Figure 5.**
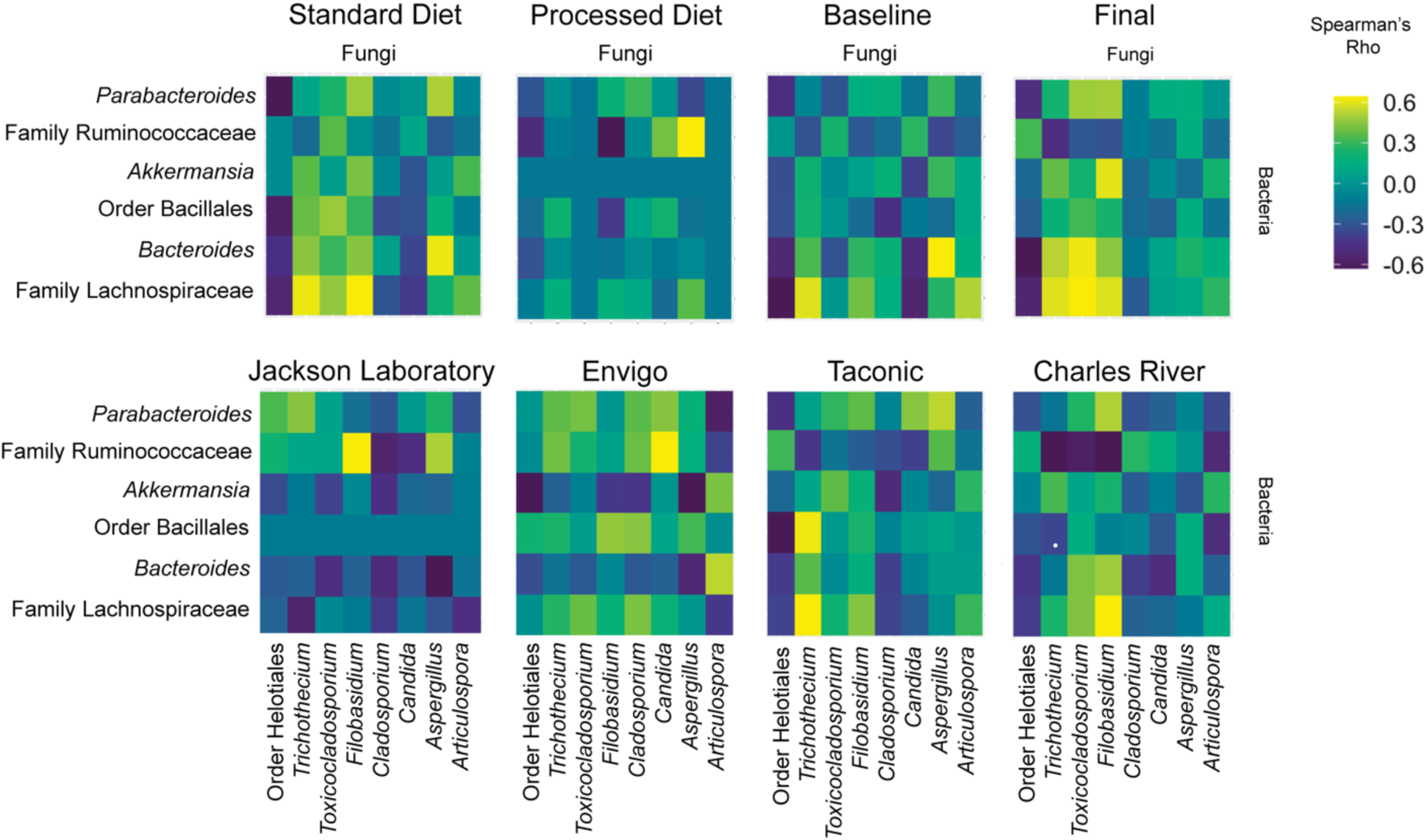
Interkingdom co-occurrence differs by vendor, diet and time. The 14 most abundant taxa are displayed (comprising > 99.2 % of variability). Color indicates strength of Bonferroni corrected Spearman’s Rho correlation coefficients. The x-axis comprises fungal taxa with bacteria on the y-axis. Complete co-occurrence networks are shown in Supplemental Figure 11.

### Gut Fungal Community Composition Correlates with Differences in Body Composition and Fat Deposition

Having established persistent differences in fungal community composition, we next asked if this change altered host metabolic tone. Because female mice are protected from diet-induced obesity,**[30]** we focused on male mice (for results from female mice, see **Supplemental Figure 10**). On processed diet, as quantified by EchoMRI and tissue collection, male mice from CR, ENV and JAX gained fat mass, but mice from TAC did not (**Figure 6B, C**). Utilizing an unbiased panel, we examined differences in serum metabolic biomarkers after a 5-hour fast. Increases in fasting leptin and ghrelin and decreases in fasting resistin appropriately correlated with increased adiposity (**Figure 6D**). While we were unable to detect lipid deposits in mice exposed to standardized diet, we noted complementary differences in hepatic steatosis (**Figure 6E**), a marker of metabolic syndrome in mice**[29]**, especially in mice from ENV. Gut bacteria can influence host gene expression in animal studies,**[29]** so we also examined gene expression in metabolically active tissues. In epidydimal white adipose tissue, differences we between vendors were noted in *PRDM16* (PR/SET domain 16, **Figure 6**). In the liver, key differences were noted in *PPARA* (Peroxisome Proliferator Activated Receptor Alpha) family, *CD36* and *GPAT1* (Glycerol-3-phosphate acyltransferase, **Figure 6H**).

**Figure 6.**
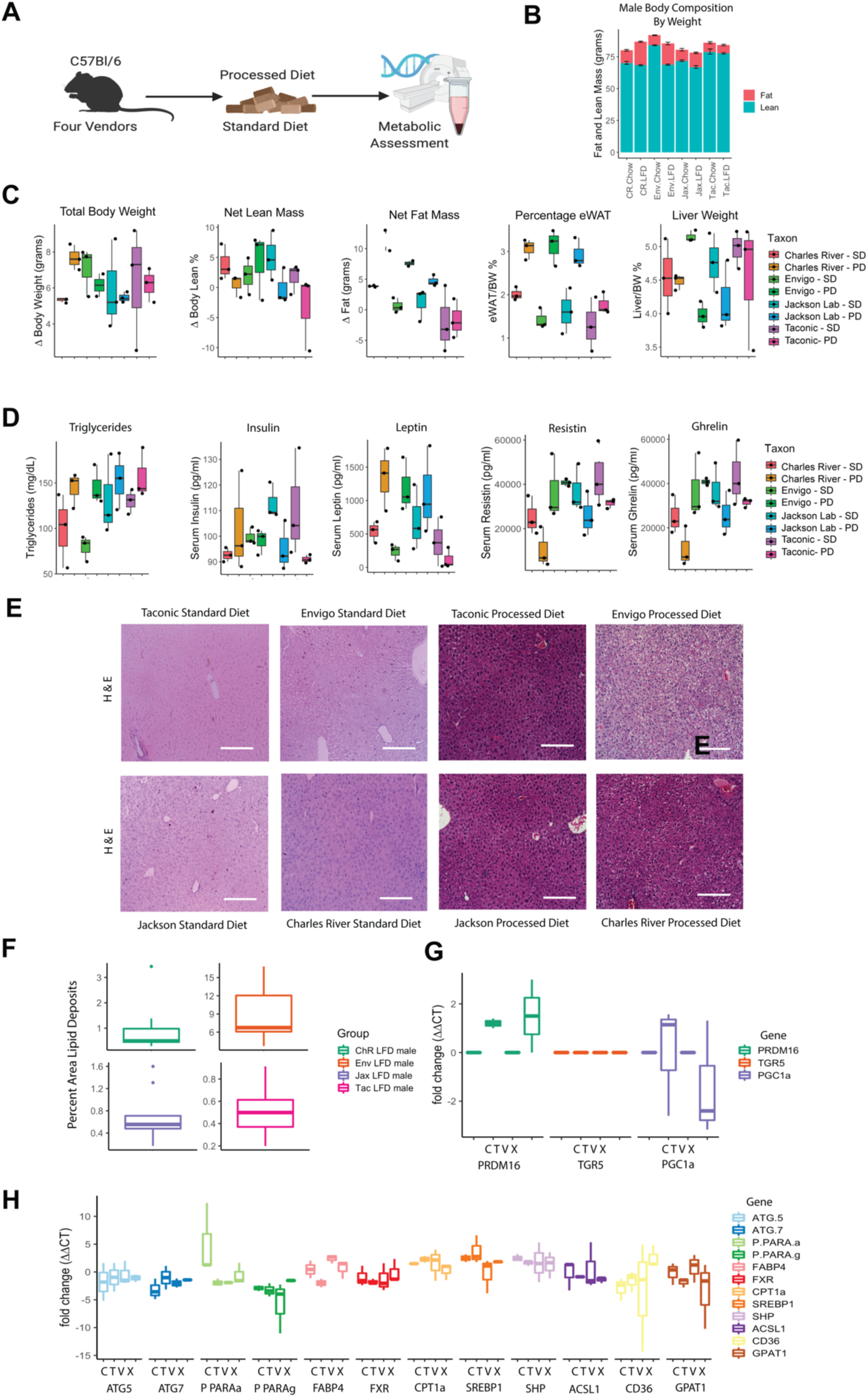
The metabolic phenotype of male mice is sensitive to gut fungal community composition. (*A*) Experimental schematic. (*B*) Male mouse body composition by EchoMRI. (*C*) Male mouse body composition by tissue collection. (*D*) Male serum metabolically active biomarkers. (*E*) Hematoxylin and eosin stained liver tissue show increased deposition after a processed diet, particularly in mice from Envigo. (*F*) Quantification of lipid deposition in males exposed to processed diet. (*G*) Gene expression in epididymal white adipose tissues. (*G*) Gene expression in liver. eWAT, epididymal white adipose tissue; PD, processed diet; SD, standard diet. Schematic illustrated using BioRender.

### Metabolic Tone is Strongly Associated with Variability in the Gut Mycobiome

To understand if the mycobiome was associated with differences in metabolic tone, we performed correlation analysis using the relative abundance of fungal genera and community diversity metrics with metabolic biomarkers (Biconjugate A-Orthogonal Residual, Spearman correlations with Bonferroni correction, **Figure 7A**). Resistin and PIA-1 (plasminogen activator inhibitor-1) were negatively correlated, while and fat mass accrual was positively correlated with the presence of several key fungi. We further explored the relationship between fungal community composition and metabolic tone by building random forest machine learning models and performing variable importance analysis to identify key fungal taxa (**Figure 7B**). We developed robust models for three metabolic outcomes that explain a sizable portion of the data variability. Differences in weight gain moderately correlated with differences in particular fungal taxa, of which the abundance and distribution of the genus *Thermomyces* was the most important. The most robust model was for serum triglyceride concentration, for which the genus *Cladosporium* was the most important taxa. Similarly, the genera *Saccharomyces* and *Aspergillus* were important for our model of fasting ghrelin concentration. Altogether, these results suggest fungal community composition is strongly associated with host metabolic tone.

**Figure 7.**
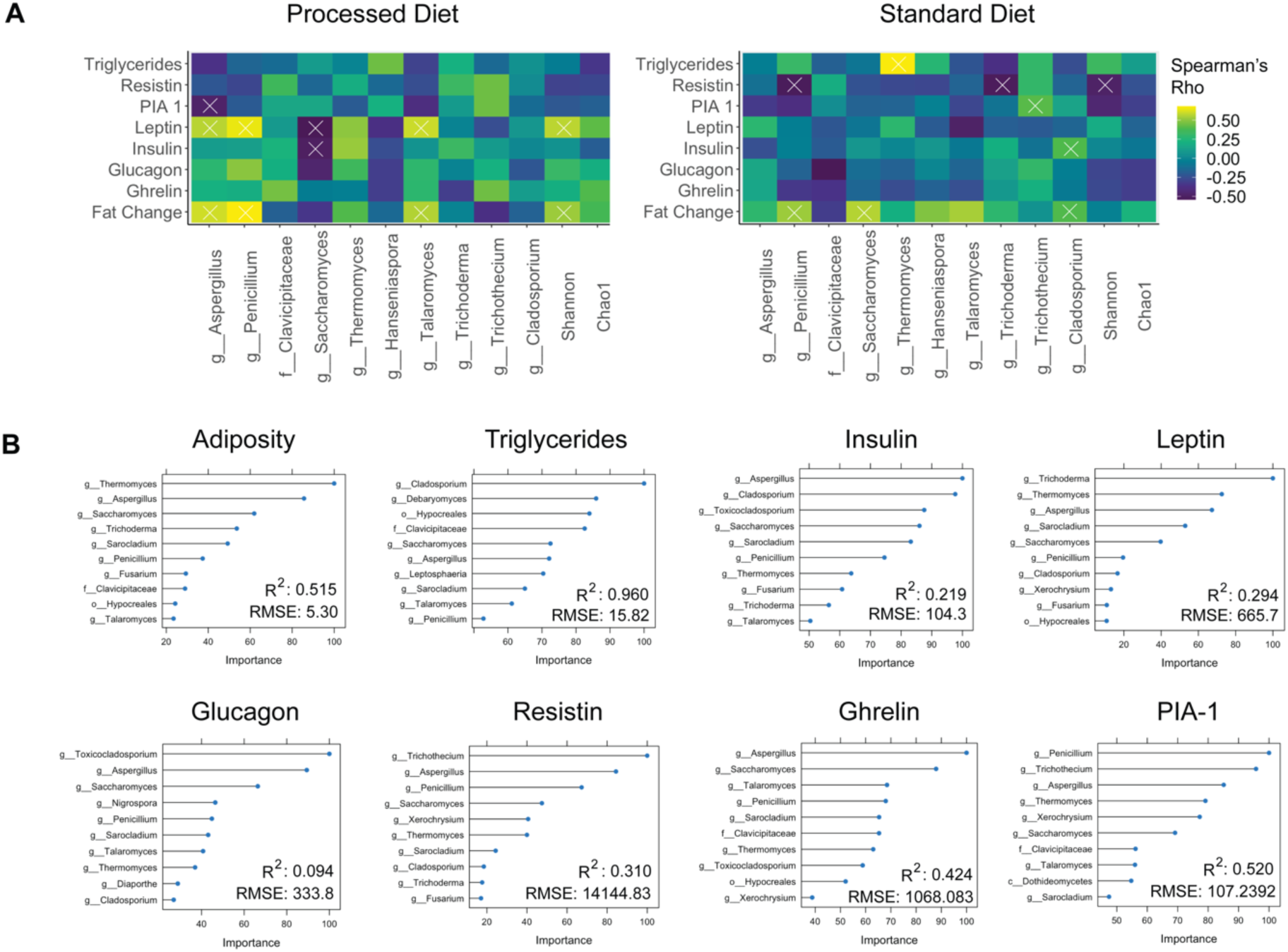
Fungal genera strongly associate with metabolic tone. (*A*) Biconjugate A-Orthogonal Residual method. X indicates p <0.05 after false discovery rate correction. (*B*) Random forest regression models showing the relative importance of a particular taxon to the model. PIA-1, plasminogen activator inhibitor-1; R^2^, correlation coefficient; RMSE, root mean square error.

## DISCUSSION

Here, we investigated whether variations in gut mycobiome abundance and composition correlate with key features of host metabolism. To address this question, we asked if differing starting mycobiomes could affect host adaptation to standardized diet and an ultra-processed diet rich in purified carbohydrates in a manner that associated with deleterious metabolic outcomes. Our key findings are the gut mycobiome of healthy mice is shaped by the environment, including diet, and significantly correlates with metabolic changes in the host. For instance, increased triglyceride concentrations and metabolic biomarkers, including deposition of hepatic lipids, correlate with increased abundance of the fungal genera *Thermomyces* and decreased *Saccharomyces*. Our results highlight the potential importance of the gut mycobiome in health and have implications for human and experimental metabolic studies.

The bacterial microbiome is strongly influenced by dietary exposure^30^ and influences host metabolism.^3^ Emerging work demonstrates dietary exposure also shapes fungal communities.^8^ However, despite evidence for fungal pattern recognition receptors in the human gut^6^ and fungal influences on disease in the human gut,^31,32^ continuous gut colonization by fungi remains controversial in humans.^33^ Convincing evidence indicates fungi colonize the mouse gut and influence host physiology^34^ and disease.^35^ Diet may have a dominant effect over host genotype on the composition of the gut bacteriome with subsequent alterations in host-microbe interactions.^36^ We observed a similar strong effect of diet on the composition of both fungal and interkingdom community composition. However, because we used mice with the same genetic background, we were able to examine the impact of the founding microbial communities mostly independent of host genomic influences. In our study, both prolonged exposure to a processed diet and length of isolation in a specific pathogen-free environment reduced fungal diversity. In turn, reduced fungal diversity was associated with increased adiposity and physiologic alterations seen in the metabolic syndrome.

In our controlled study, we found key differences in baseline mycobiome composition reflect differences in metabolic tone in response to diet. For metabolic studies in mice, the choice of vendor, shipment, diet and housing may play instrumental roles in shaping outcomes, which should encourage further caution in drawing causative relationships. Validating outcomes in mice from several vendors or across multiple shipments may be necessary to address this potential confounder. The implication for human microbiome studies, which often examine only bacteria and sample only fecal communities, is that the mycobiome may have unappreciated effects on microbiome-associated outcomes.

Exposure to high-fat diet may alter fungal and interkingdom community composition.^8^ Our work suggests complex alterations in co-abundance networks are associated with diet. Fungal cell wall components are a major point of interaction between fungi and bacteria in the environment.^6^ For example, the abundance of a major gut bacteria, *Bacteroides thetaiotaomicron*, can be influenced by the presence of mannan in fungal cell walls.^15^ Similarly, fungal chitin influences the composition of anaerobic bacteria.^37^ Our co-occurrence correlation analysis suggests the loss of key fungi during dietary exposure was closely related to differences in bacterial community composition, which may represent niche replacement in the face of a shifting dietary environment or disruption of interkingdom metabolomic networks. This study extends previous work^8^ in two important avenues. First, we examined the impact of an ultra-processed diet on gut communities. This approach allowed us to identify more subtle physiologic changes in host metabolic tone. Second, by using mice from different vendors, we extended these observations by observing the effect of different founding mycobiomes on host metabolism.

Among the myriad metabolites and effectors that likely connect gut microbial communities to metabolism, the gut bacteriome may influence host metabolism through several major mechanisms.^38^ While classically attributed to bacteria, fungi play an often underappreciated role in the production of metabolites and may interact with host physiology via analogous mechanisms.

One of the best-known fungal pattern recognition receptors in humans is Dectin-1, which recognizes fungal β-glucan. In addition to immune cells,^6^ adipose tissue expresses Dectin-1.^39^ In humans, obesity is associated with increased Dectin-1 expression in adipose tissue.^39^ In mice, Dectin-1 plays a MyD88-independent role in diet-induced obesity.^39^ In MyD88-deficient adipose tissue, Dectin-1 was upregulated in both adipocytes and adipose-associated macrophages. Furthermore, blockade of Dectin-1 led to improved glucose sensitivity and decreased numbers of CD11c^+^ macrophages. The reverse was also true in Dectin-1 activation.^39^ Thus, mycobiome-driven antigen presenting cell education in the gut could directly influence systemic metabolic tone.

Another important function of gut microbes, often attributed solely to bacteria, is the production of secondary bile acids (BAs). Our understanding role of BAs has evolved from simple detergents, to hormones intimately related to multiple metabolic processes, revealing important roles of BAs in dyslipidemia and type 2 diabetes.^40^ BAs are a significant source of host-microbiome interaction via cellular receptors such as TGR5^41^ and FXR.^40^ Therefore, BA composition is interrelated with metabolic hormones such as leptin,^42,43^ resistin,^44^ ghrelin, GLP-1 (glucose-like peptide-1) and peptide YY.^45^ However, fungi also produce BAs and likely participate in the gut BA pool to an underappreciated extent.^46^ *Fusarium*, which we found was key to the structure of processed diet-induced weight gain-resistant TAC mice, is a prolific metabolizer of deoxycholic acid. Other key fungi, such as *Aspergillus* and *Penicillium*, can also produce secondary BAs.^46^ BAs also influence the stability of other fungal metabolites. For example, luminal BAs^47^ influence the stability of a prominent lipase in *Thermomyces*, which we observed as the taxa most significantly associated with weight gain on a processed diet. Intriguingly, in our model, these differences correlated with differences in metabolic hormones, including leptin, resistin and ghrelin. These findings may suggest a potential role for fungi in host metabolic processes via BA signaling.

In summary, these data indicate the gut mycobiome in healthy mice is highly variable and responds to disturbances such as changes in the environment and diet. Despite these ecological pressures, resilient differences in gut mycobiome composition in healthy mice strongly associate with differences in host metabolic tone, including differential fat deposition, metabolic biomarkers and gene expression in metabolic tissues. We also highlighted two fungal-derived products with plausible effects on host metabolism. While there are potentially thousands of metabolically active fungal products, our work argues for future work in defined gnotobiotic animal models to further elucidate the mechanisms of mycobiome-host interaction. Finally, our findings suggest that differences in the gut mycobiome may be an underappreciated source of variability in health and metabolic outcomes in response to dietary interventions.

## Acknowledgments

The authors would like to thank the Children’s Foundation Research Institute at Le Bonheur Children’s Hospital for providing scientific editing.

## Data Availability Statement

Sequences are available via the National Center for Biotechnology Information Short Read Archive (BioProject PRJNA597168). Operational taxonomic unit, metadata tables, univariate analysis results and R code are available from github.com/WillisLungLab/Vendors_gut_mycobiome.

