## Supplemental File for "The gut mycobiome of healthy mice is shaped by the environment and shapes metabolic outcomes in response to diet"

**Graphical Abstract:**

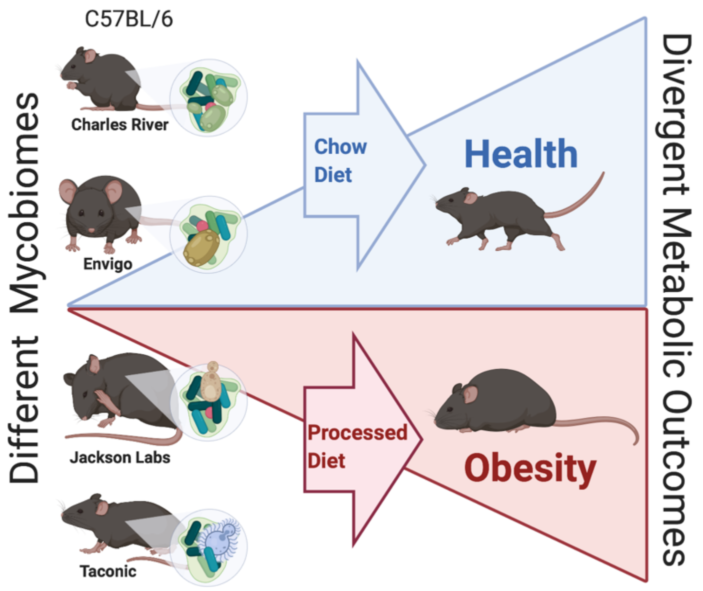


**Supplemental Methods:**

*Animals:*

All procedures involving animals were performed in accordance with the National Institute of Health Guidelines for the Care and Use of Experimental animals and were approved by the University of Tennessee Health Science Center Institutional Animal Care and Use Committee. Animals were housed in sterile microisolators upon arrival in the same room under a standard 12-hour light-dark cycle.

*Diets:*

| Dietary Components | Chow Diet (Envigo 7912) | Processed Diet (Research Diets D D12450B) |
|---|---|---|
| Protein | 19.1% Kcal | 20% Kcal |
| Fat | 5.8% Kcal | 10% Kcal |
| Carbohydrate | 44.3% Kcal | 70% Kcal |
| Fiber | 4.6% | - |
| Neutral Detergent Fiber | 13.7% | - |
| Minerals | 4% | 4.74% |
| Energy Density | 3.1 Kcal/g | 2.82 Kcal/g |

*Tissue collection:*
At the experimental endpoint, inguinal adipose depots (iWAT), gonadal adipose depots (gWAT), intrascapular brown adipose tissue, (iBAT), and liver were immediately dissected and cleared of any connective tissue before being weighed to determine specific organ/tissue weights. Tissues were then snap frozen in liquid nitrogen or fixed in 10% neutral buffered formalin for histology.

*Serum biochemistry:*
Freshly obtained blood samples were allowed to clot for 30 minutes at room temperature before undergoing centrifugation at 5,000 rcf for 5 minutes to obtain serum samples.

*Serum metabolic biomarkers:*
We utilized an 8-plex Bio-Plex Pro Mouse Diabetes Panel from Bio-Rad (cat. #171F7001M, Hercules, CA) to quantify ghrelin, GIP, GLP-1, glucagon, insulin, leptin, PAI-1 and resistin in mouse serum, according to the manufacturer's instructions.

*Body Composition:*
Body composition (fat and fat-free mass) were measured non-invasively every other week using an EchoMRI 1100 system (EchoMRI, Houston, TX). System calibrations were performed before each session.

*Histology:*
Liver was harvested and fixed overnight in 4% paraformaldehyde. Tissues were processed (Tissue-Tek V.I.P, Sakura Finetek, Torrance, CA) and embedded in paraffin. For representative images, samples were cut (5.0 µm) and placed on adhesive coated slides (Newcomer Supply, Madison, WI), deparaffinized, rehydrated and H&E stained (Cat. No. H-3502, Vector Laboratories, Burlington, Canada). For pathology assessment of lipid deposition, samples were imaged in triplicate for representative images.

*PCR:*
Tissues were collected in TRIzol reagent (Ambion, Austin, TX). RNA was isolated using the TRIzol and chloroform method, and RNA purity was validated through UV-Vis spectrophotometry using a Nanodrop Lite (Thermo Scientific, Wilmington, DE). Total RNA (1.0 µg) was reverse-transcribed to complementary DNA (cDNA) using a Transcriptor First Strand cDNA Synthesis Kit (Roche, Indianapolis, IN) according to the manufacturer's instructions. RT-PCR amplification comprised an initial denaturation step (95°C for 10 min), 45 cycles of denaturation (95°C for 10s), annealing (55°C for 20s) and extension (60°C for 30s), followed by a final incubation at 55°C for 30s and cooling at 40°C for 30s. All measurements were normalized by the expression of the GAPDH gene, a stable housekeeping gene. Gene expression was determined using the delta-delta Ct method: $2^{-\Delta\Delta CT}$ ($^{\Delta\Delta}$CT = [Ct (target gene) - $Ct(GAPDH)]_{tested}$ - [Ct(target gene) -$Ct(GAPDH)]_{control}$) and displayed as relative mRNA levels.

*DNA extraction and Illumina MiSeq sequencing:*
Murine jejunal and colonic luminal material samples were resuspended in 500 mL TNES buffer containing 200 units lyticase and 100 mL 0.1/0.5 (50/50 Vol.) zirconia beads. Incubation was performed for 20 minutes at 37°C. Following mechanical disruption using ultra-high-speed bead beating, 20 mg proteinase K was added to all samples, which were incubated overnight at 55°C with agitation. Total

DNA was extracted using chloroform isoamyl alcohol, and total DNA concentration per mg stool was determined by qRT-PCR. Purified DNA samples were sent to the Argonne National Laboratory (Lemont, IL) for amplicon sequencing using the NextGen Illumina MiSeq platform, utilizing 16S rRNA MiSeq for bacteria and archaea and parallel ITS2 rDNA sequencing for fungi. Blank samples for the jejunal sequencing run passed through the entire collection, extraction and amplification process remained free of DNA amplification. Fecal samples were excluded from downstream analysis due to detection of a suspected contaminant apparently introduced during the MiSeq library preparation.

*Bioinformatics:*
Sequencing data were processed using QIIME 1.9.1. Sequences were demultiplexed, denoised and clustered into operational taxonomic units (OTUs). For bacteria, sequences were aligned via PyNAST, and taxonomy was assigned against the SILVA database. For fungi, sequences were aligned, and taxonomy was assigned using the UNITE (dynamic setting) database. All OTU count data followed a negative-binomial distribution. Processed data were then imported into Calypso 8.84 for further data analysis and visualization. Additional data analysis was performed in R. On import in Calypso, all mitochondrial sequences were discarded. For bacteria, any samples with less than 100 sequence reads were discarded, resulting in the removal of 0 samples from downstream analysis. For fungi, samples with less than 100 sequence reads were discarded, resulting in the removal of 21 presumed blank samples from downstream analysis. Processed data underwent a Hellinger transformation (square root of total sum normalization). We then utilized principal components analysis (PCA), principal coordinates analysis (PCoA) and canonical correspondence analysis (CCA) plots of Bray-Curtis dissimilarity distances to visualize beta diversity. Statistical significance of beta diversity clustering was then assessed using CCA and permutational multivariate analysis of variance (PERMANOVA) followed by permutational analysis of multivariate dispersions (PERMDISP2) to assess the homogeneity of group variance (distance to centroid). To assess alpha diversity, we rarified bacterial samples to a depth of 4642 reads and rarefied fungal samples to a depth of 889 reads. Then, the Shannon and Chao1 diversity indices were calculated and differences assessed using ANOVA. On unrarefied data, we also performed univariate analyses using analysis of comparison of microbiomes (ANCOM), core microbiome analysis, supervised partial least squares discriminant analysis (sPLS-DA), mixed-effect regression with sex as a random effect, and negative binomial regression (*DESeq2* function) to quantify the differences in the relative abundance of specific microbial taxa, as appropriate. We used linear discriminant analysis of effect size (LEfSe) to perform high-dimensional biomarker identification. Co-occurrence networks using the top 15 most abundant taxa (representing 99.45 % ± 1.3% of the interkingdom communities) were calculated using pairwise Spearman correlations with Bonferroni correction of taxon counts of which all were significant.

*Modeling:*
Generalized linear modeling at the multivariate and univariate (with Bonferroni correction) levels using a negative-binomial distribution with likelihood ratio tests and resampled p-values to calculate significance was performed in R utilizing the *mvabund* package. Bacterial and Fungal OTUs were modeled separately to test the effects of vendor, diet, sex, and time of measurement of the microbiome and identify the taxa that significantly differ by variable.

Random forest regression (RFR) was used to regress changes in serology to variation in taxonomic interkingdom community composition. For this model, Hellinger transformed OTU counts were separated into 60% training and 40% validation sets with 10-fold cross-validations to test model accuracy. Data were preprocessed by removing zero and near zero features, scaling and centering all features, and removing highly correlated features (rho > 0.90). Feature selection to identify the most important taxa in the random forest regression was conducted using the *varimp* (*caret*) function where a LOESS (locally estimated scatterplot smoothing) function is fit between the outcome and the predictor. We assigned an importance cutoff at 10 as the point when adding in new variables becomes asymptotic for all models (additional taxa do not increase model accuracy meaningfully). An $R^2$ statistic was calculated for this model against the null model intercept. This number was returned as a relative measure of variable importance. Error was assessed by calculating the root mean standard error (RMSE), the average difference between the observed known values of the outcome and the predicted value by the model. This model was constructed and repeated for changes in fat, triglycerides, insulin, leptin, glucagon, resistin, ghrelin, and PIA1 levels measured in the mice.

Supplemental Figures

Supplemental Figure 1.

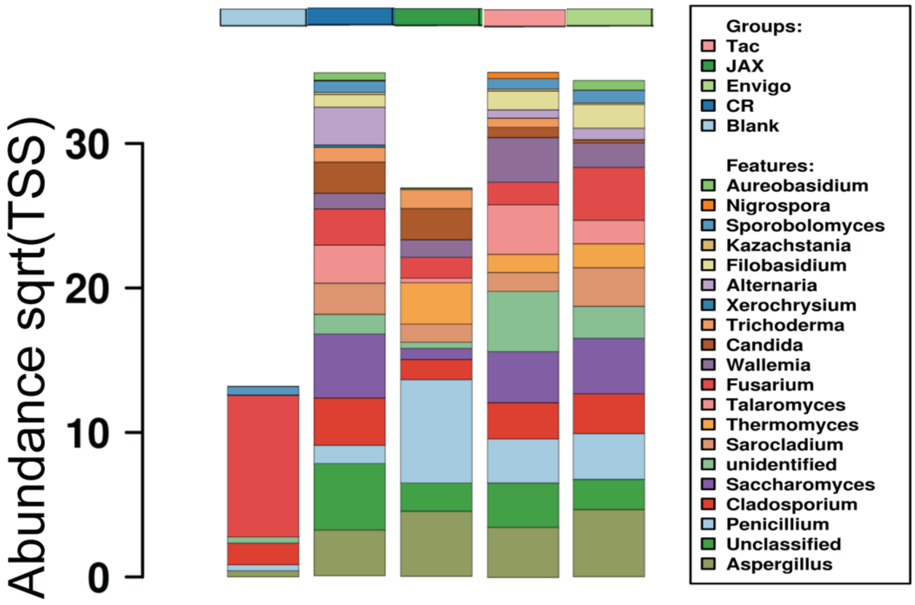


**Supplemental Figure 1.** Sequence and process controls show now consistent patterns of contamination. Pooled bar charts at the genus level. Fusarium is the most frequent contaminant detected.

Supplemental Figure 2.

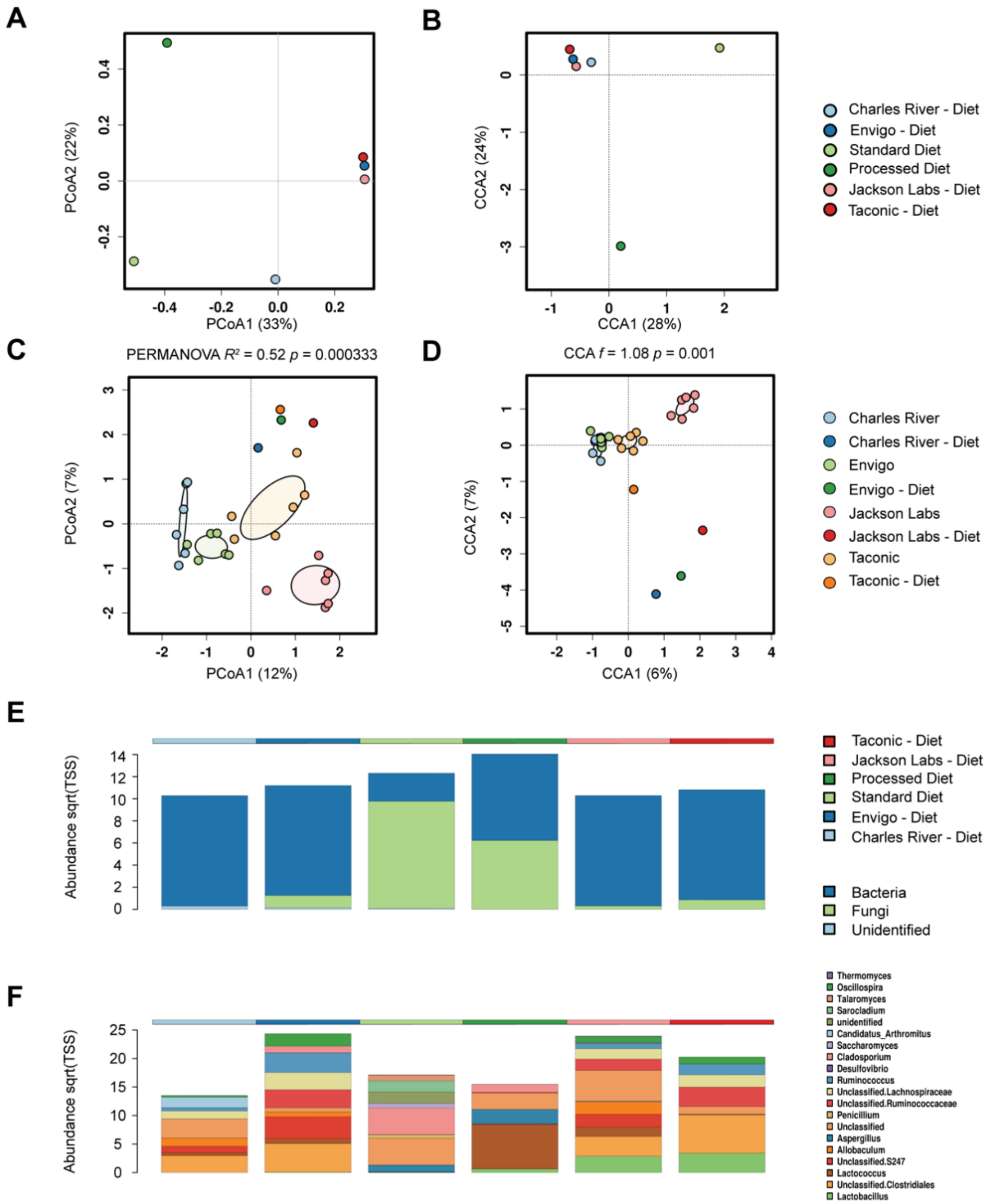


**Supplemental Figure 2.** Multi-kingdom composition of mouse diets.

Supplemental Figure 3.

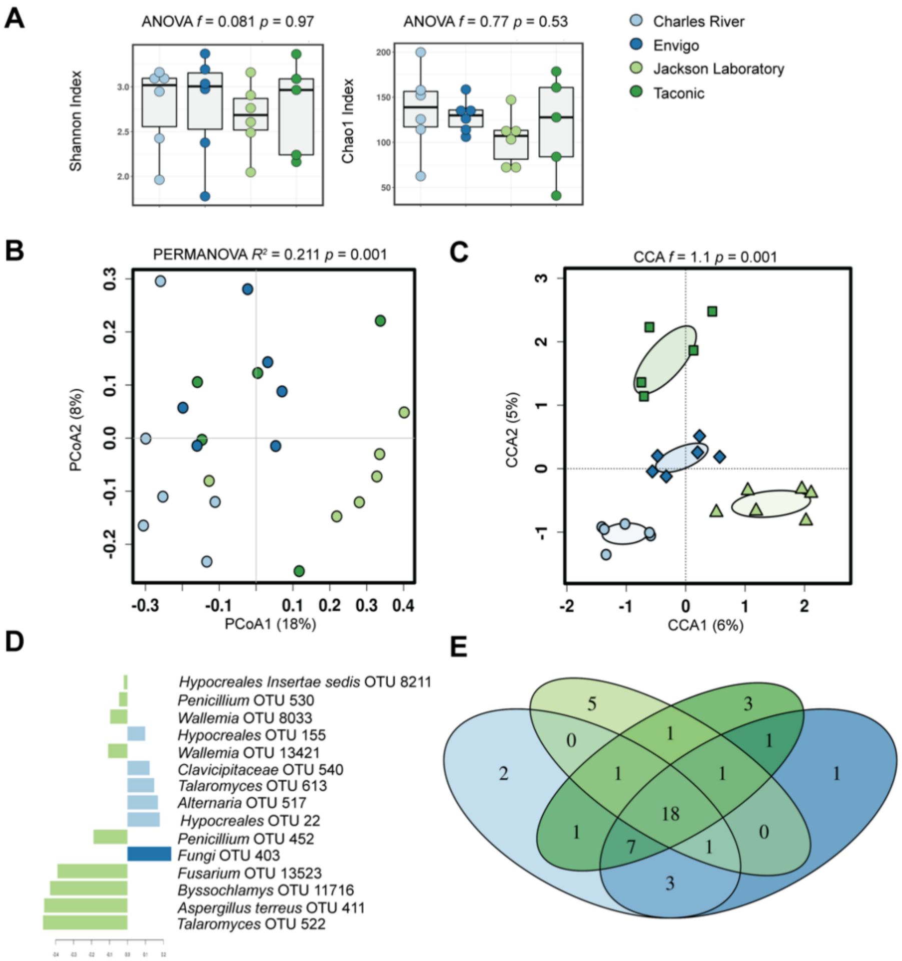


**Supplemental Figure 3.** The baseline fungal community composition differs between vendors.

Supplemental Figure 4.

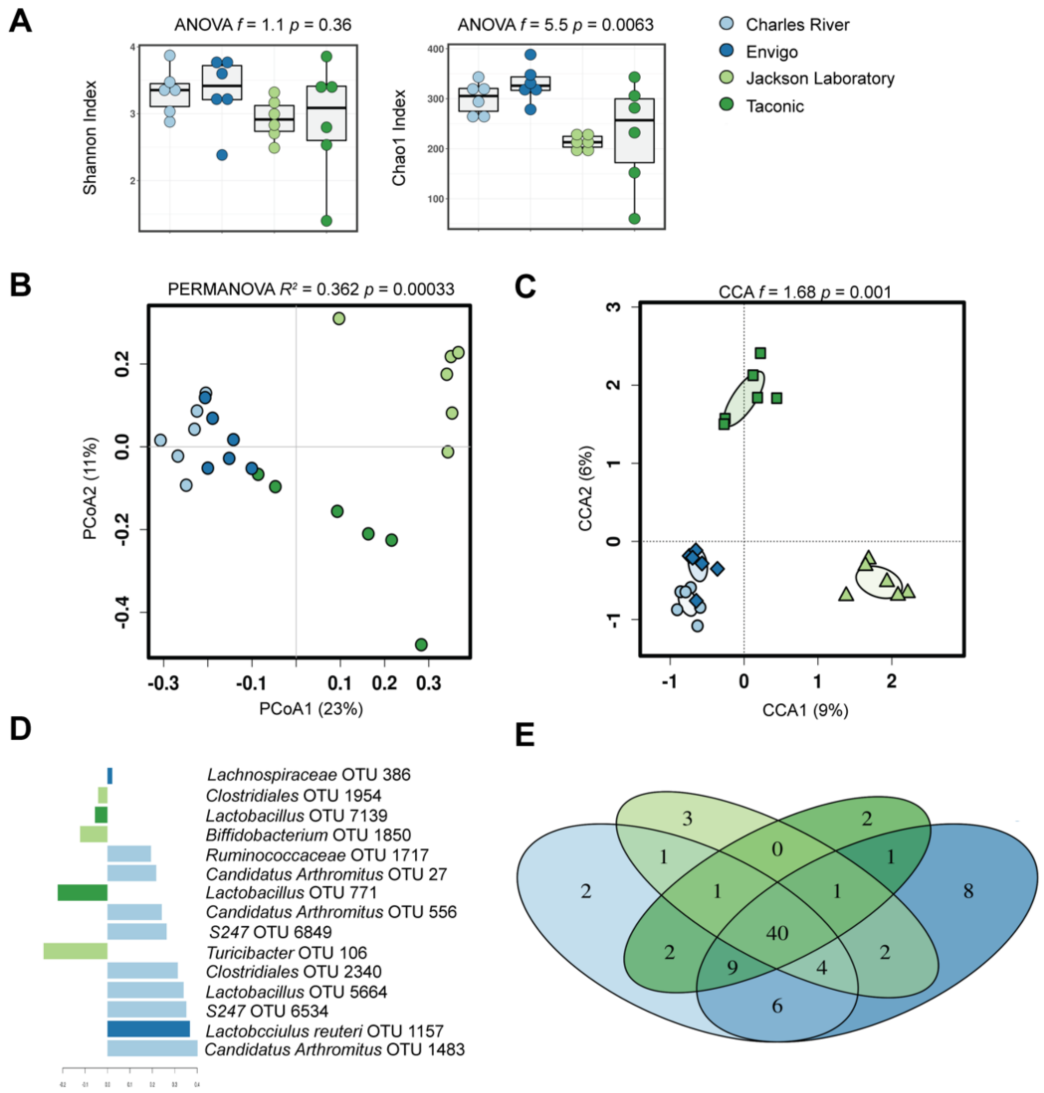


**Supplemental Figure 4.** The baseline interkingdom community composition differs between vendors.

Supplemental Figure 5.

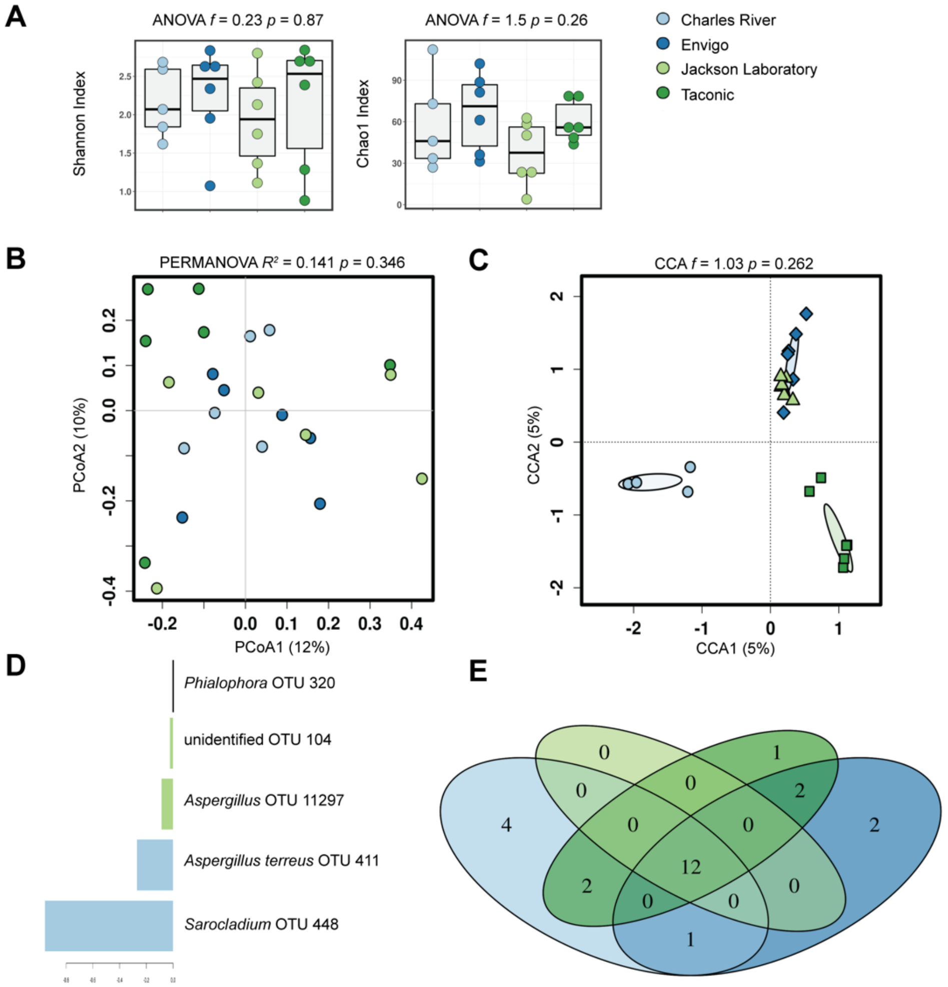


**Supplemental Figure 5.** Fungal community composition after 8-week exposure to standardized diet.

Supplemental Figure 6

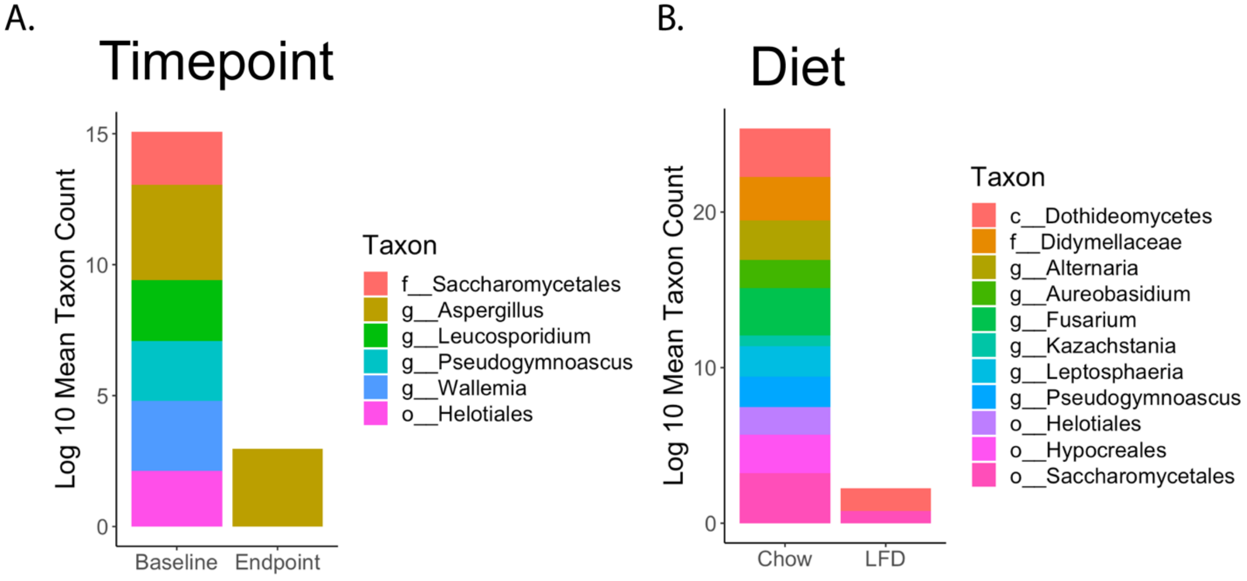


**Supplemental Figure 6**. Stacked bar plots of significant univariate MVABUND modeling of taxa responses. A. $Log_{10}$ of mean taxon abundance separated by baseline and endpoint measurements. B. $Log_{10}$ of mean taxon abundance separated by diet.

Supplemental Figure 7.

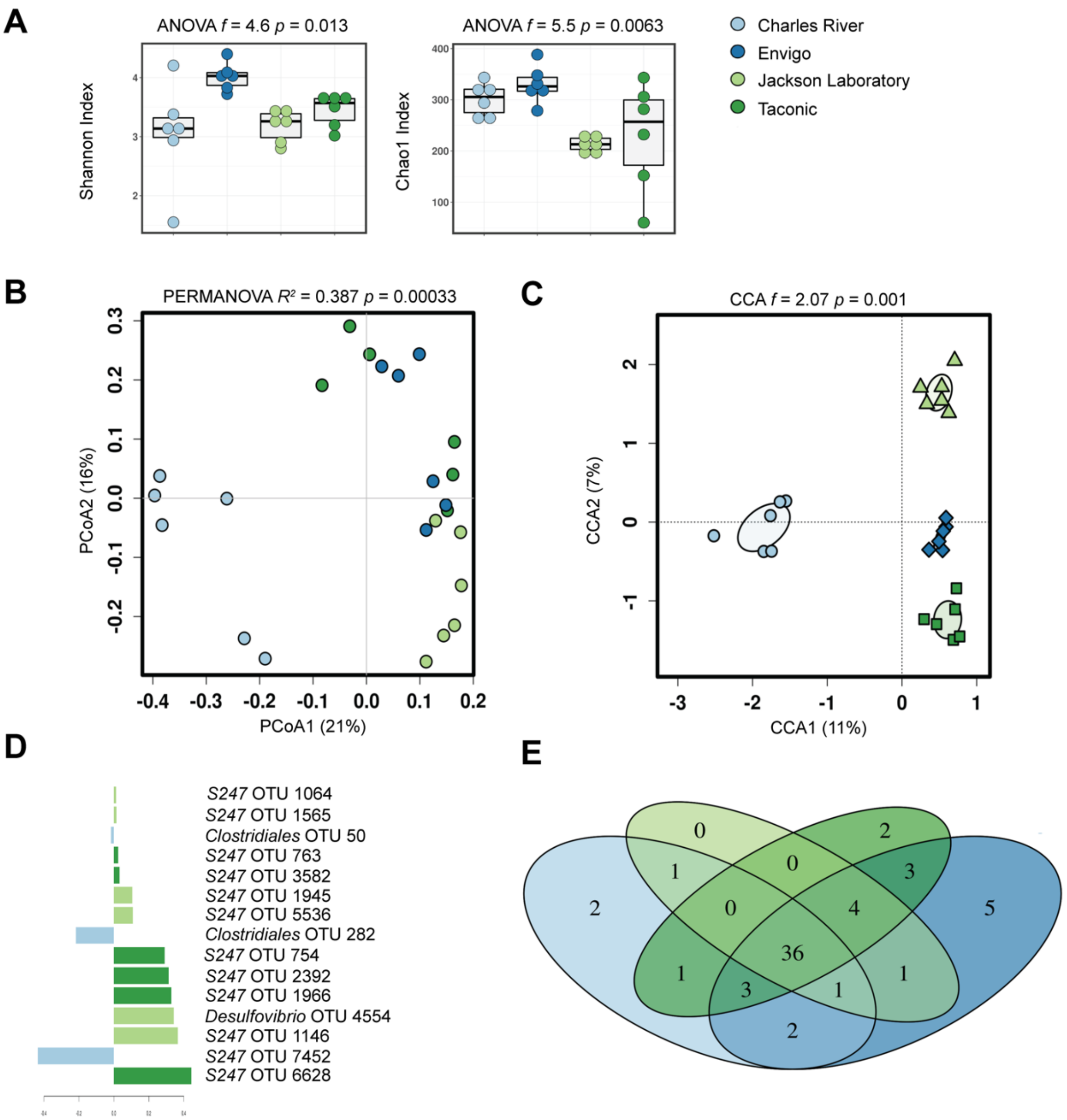


**Supplemental Figure 7.** Interkingdom community composition differs after exposure to standardized diet.

Supplemental Figure 8.

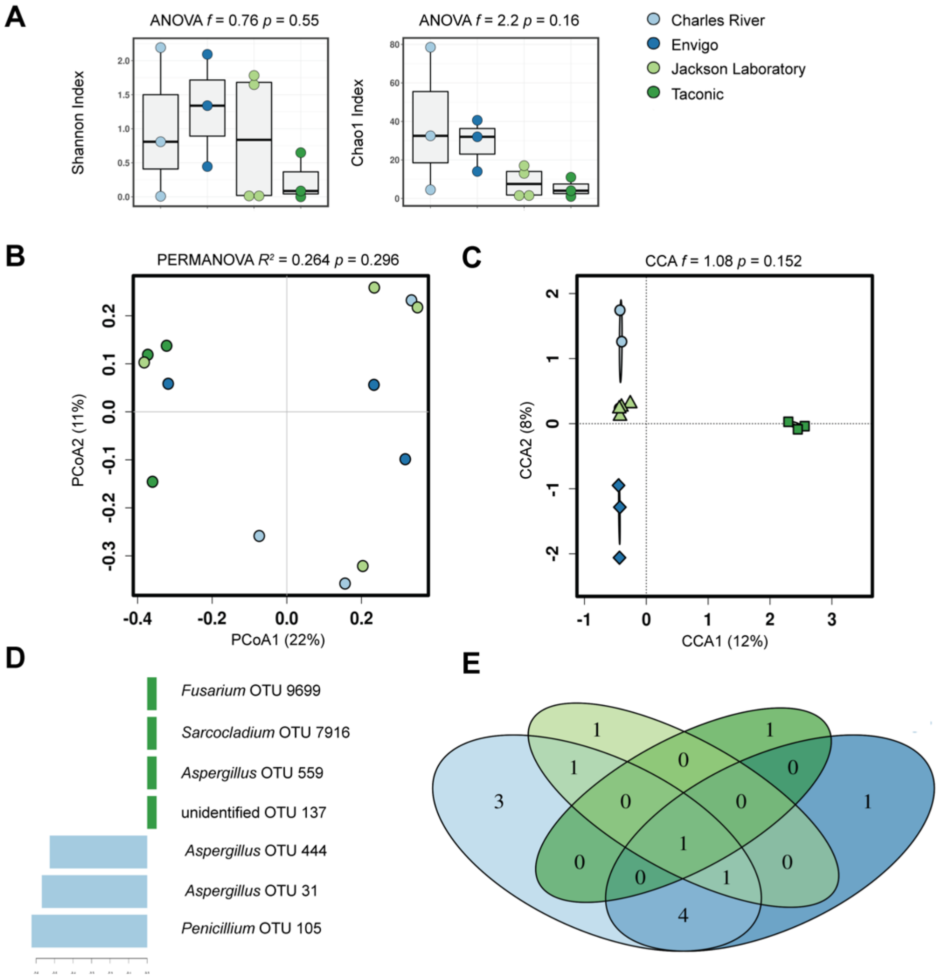


**Supplemental Figure 8.** Fungal community composition after 8-week exposure to processed diet.

Supplemental Figure 9.

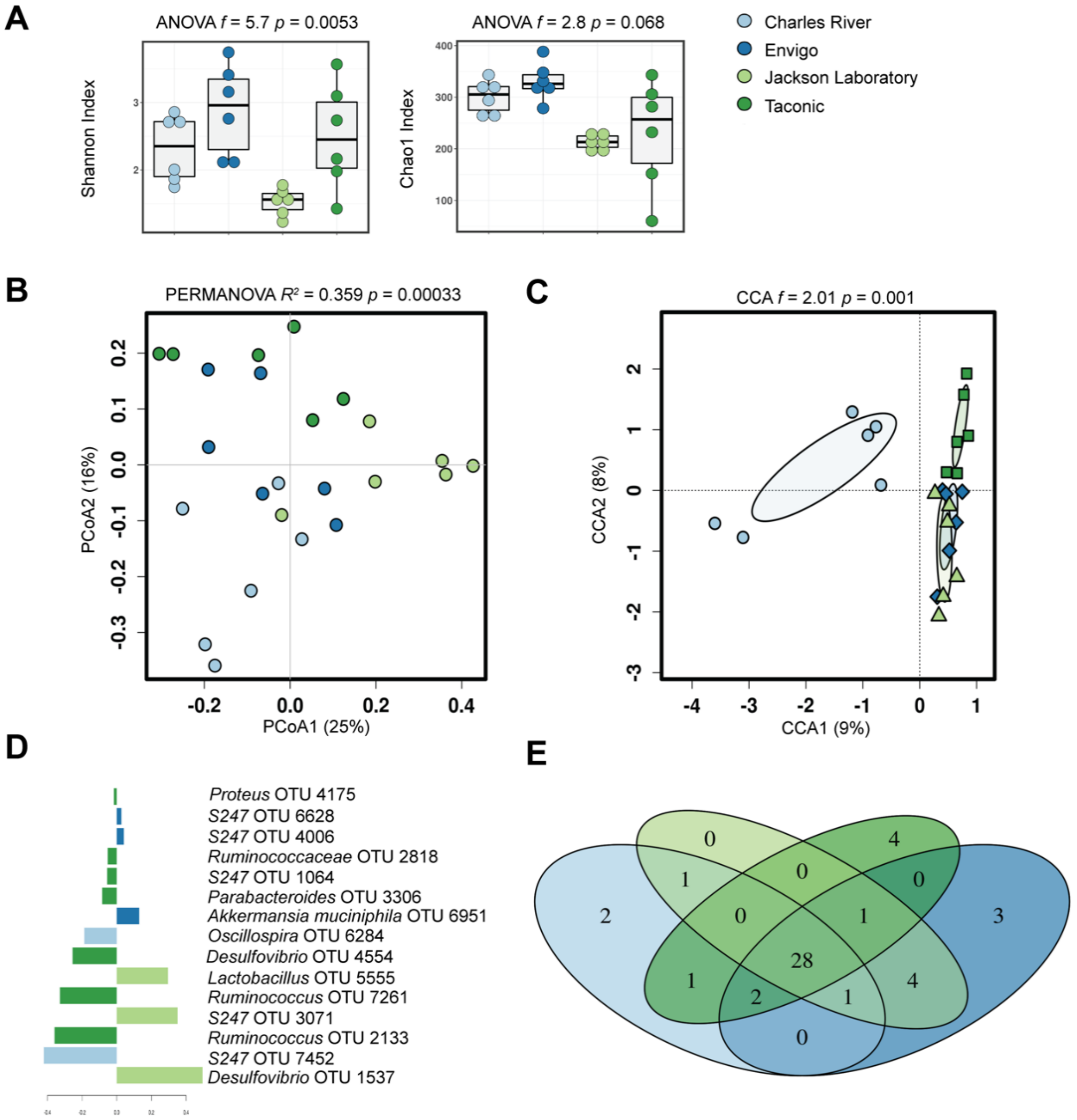


**Supplemental Figure 9.** The baseline interkingdom community composition differs after exposure to processed diet.

Supplemental Figure 10.

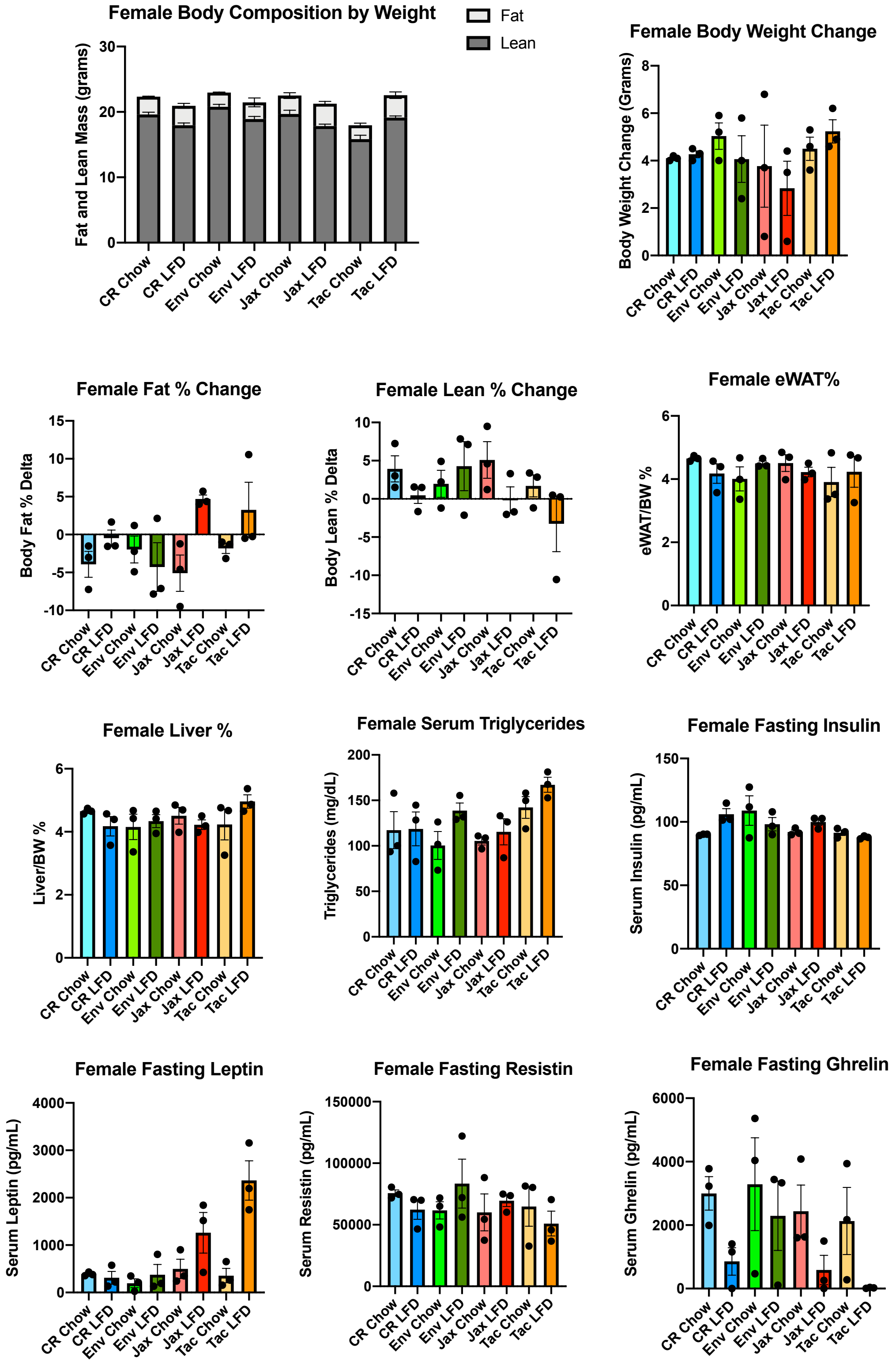


**Supplemental Figure 10.** Metabolic tone of female mice.

Supplemental Figure 11.

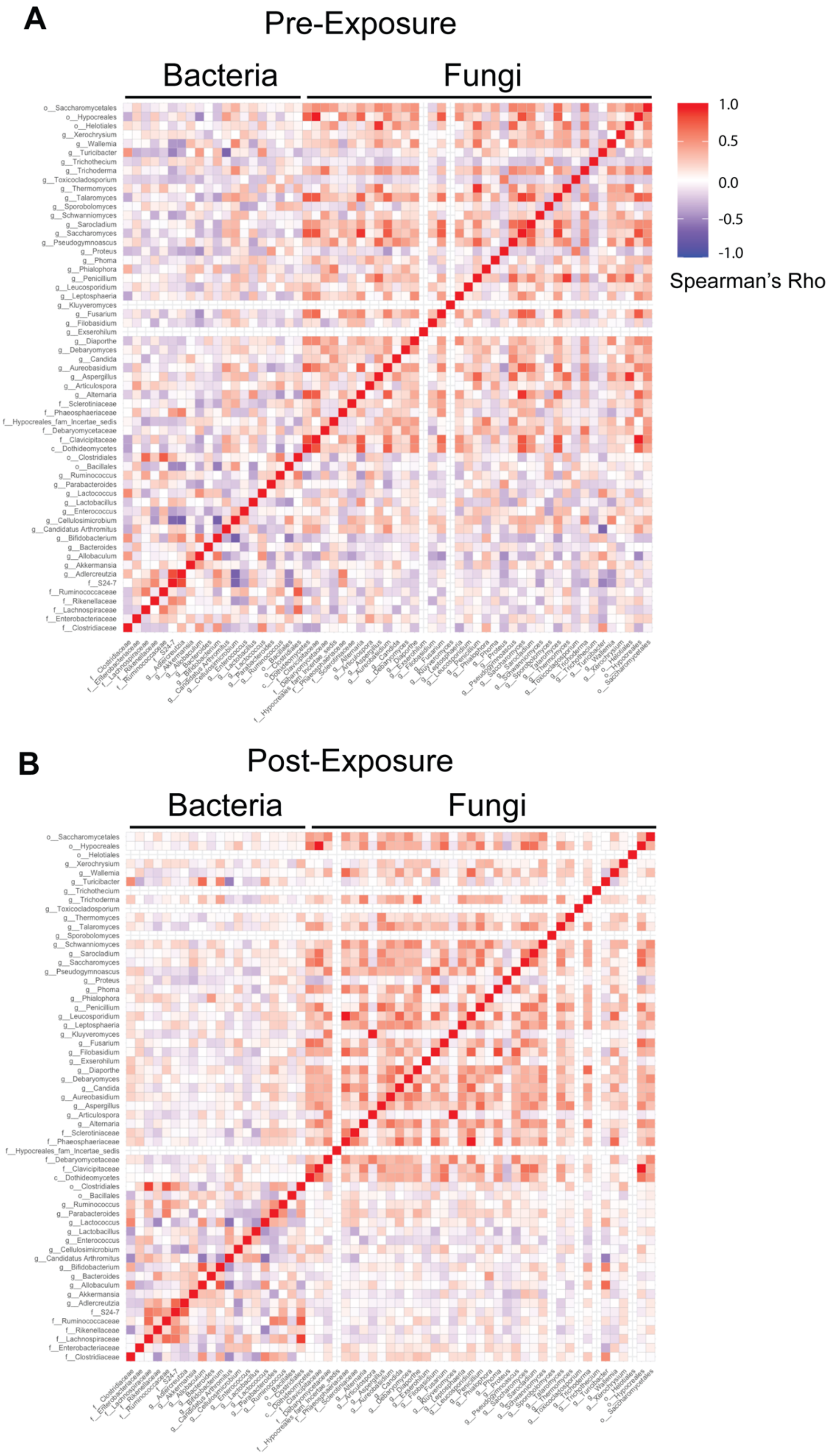


**Figure 11.** Heatmap of complete interkingdom variation before and after dietary exposure.
